# Effective Public Apologies

**DOI:** 10.1101/564880

**Authors:** Hoh Kim, Jerald D. Kralik, Kyongsik Yun, Yong-an Chung, Jaeseung Jeong

**Author notes:** **Address correspondence:** Jaeseung Jeong, Ph.D. and Jerald D. Kralik, Ph.D., Department of Bio and Brain Engineering, Korea Advanced Institute of Science and Technology (KAIST), 373-1 Kuseong-dong, Yuseong-gu, Daejeon, South Korea 34141; : and (e-mail).

## Abstract

Communicating with the public after corporate crises is often necessary, yet little evidence provides guidance. To address this, our theoretical and content analyses of public apologies revealed 12 key content elements. From these, we developed a *basic apology*, and tested its effectiveness alone, and with additional content. In two experiments involving river contamination, the *basic apology* was effective and improved with additional content. In Experiment 1, effectiveness involved *actions taken* to reduce harm and reoccurrence. Experiment 2 increased the hazard to carcinogenic chemicals, and one apology was superior: the *basic apology* plus statements of *recovery* efforts and *defense* of company actions. The two experiments show that crisis severity influences apology effectiveness. Experiment 3 found that clarifying *causality* helps convince people that the crisis source is identified and the problem resolved. Our findings show that an optimal public apology is comprehensive, and details the causes and actions taken to prevent reoccurrence.

## Introduction

When corporations are involved in crisis events, they must determine whether to issue a public statement, and if so, what to say. Among other things, a company’s reputation and thus future success is at risk, ultimately requiring forgiveness for their role in the crisis. Often this requires some form of apology, but what form should it take? Recent incidences such as a doctor being violently dragged from a United Airlines flight, followed by ineffective attempts to assuage public outrage, vividly demonstrates the significance of public statements and how an improper public apology can do more harm than good (McCann, 2017). Yet there is surprisingly little empirical evidence for public apology effectiveness, with many corporations and their advisors believing that too much information may be more harmful than not (Kim, Park, Cha & Jeong, 2015; Page, 2014; Philpot & Hornsey, 2008; Shaw, Wild & Colquitt, 2003). Moreover, the evidence thus far is mixed, warranting further examination (Coombs & Holladay, 2008; Kim et al., 2015; Philpot & Hornsey, 2008).

Although interpersonal conflict could shed light on public apologies, reaching a consensus on the critical components remains elusive, with multiple frameworks of overlapping and unique features (Battistella, 2014; Blum-Kulka & Olshtain, 1984; Lewicki & Polin, 2012; Lewicki, Polin & Lount, 2016; Scher & Darley, 1997; Schumann, 2014). Moreover, whether interpersonal conflict results apply to public apologies after corporate crises also is unknown. In fact, when one study compared interpersonal versus intergroup apologies directly, evidence for effectiveness was found only for the interpersonal case, thus calling into question the generalizability of the interpersonal findings for public apologies (Philpot & Hornsey, 2008). Therefore, we first undertook both theoretical and content analyses of public apologies, which we then used to generate the current experimental hypotheses.

Our theoretical analysis of public apologies builds from the model developed in Kim et al. (2015), and focuses on the underlying social-psychological-neurobiological principles that underpin reaction to crisis events and subsequent statements about them: i.e., victim harm and reparations for it, fairness, justice, and future threat (Kralik, 2017; Lee et al., 2018a; Lee et al., 2018b). To address these fundamental concerns, we predict that the key components of a public apology should be (1) acknowledgement of the event and explanation of what occurred; (2) acknowledgement of harm, including (a) its severity, and (b) a gesture of empathy (demonstrating actual understanding) such as regret, remorse, or repentance; (3) why it occurred, in terms of actual causal factors that led to it and who was involved; (4) assignment of responsibility (to properly redress wrongs); and evidence for (5) adequate redress of adverse consequences (such as repair or recovery efforts), and (6) forbearance (i.e., describing how the causal factor is contained and potential reoccurrence unlikely).

In fact, substantial empirical evidence in the interpersonal domain supports effectiveness of each of these message components (Anderson, Linden & Habra, 2006; Boothman, Blackwell, Campbell Jr, Commiskey & Anderson, 2009; Frantz & Bennigson, 2005; Tomlinson, Dineen & Lewicki, 2004; Tucker, Turner, Barling, Reid & Elving, 2006; Zechmeister, Garcia, Romero & Vas, 2004). Moreover, it is becoming clearer that more thorough apologies appear to be the most effective (Kirchhoff, Wagner & Strack, 2012; Lewicki & Polin, 2012; Lewicki et al., 2016; Scher & Darley, 1997; Schumann, 2014). However, whether similar, more comprehensive statements should be used in public statements remains unclear. Indeed, it appears that companies tend to minimize what is stated publicly (Page, 2014; Shaw et al., 2003).

To be sure, real-world crises may involve factors that yet escape us, and so it is also valuable to examine solutions derived in the field. We therefore also conducted a content analysis of public statements actually issued by corporations in response to major crises in South Korea (approximately 100 cases) and the United States (approximately 30 cases) from the period of 1980s-2012 via news sources (e.g., newspaper, magazine, television) and crisis case studies in journals and public relations and crisis management textbooks. Our analysis of their contents generated 12 content types listed in Table 1.

**Table 1.**
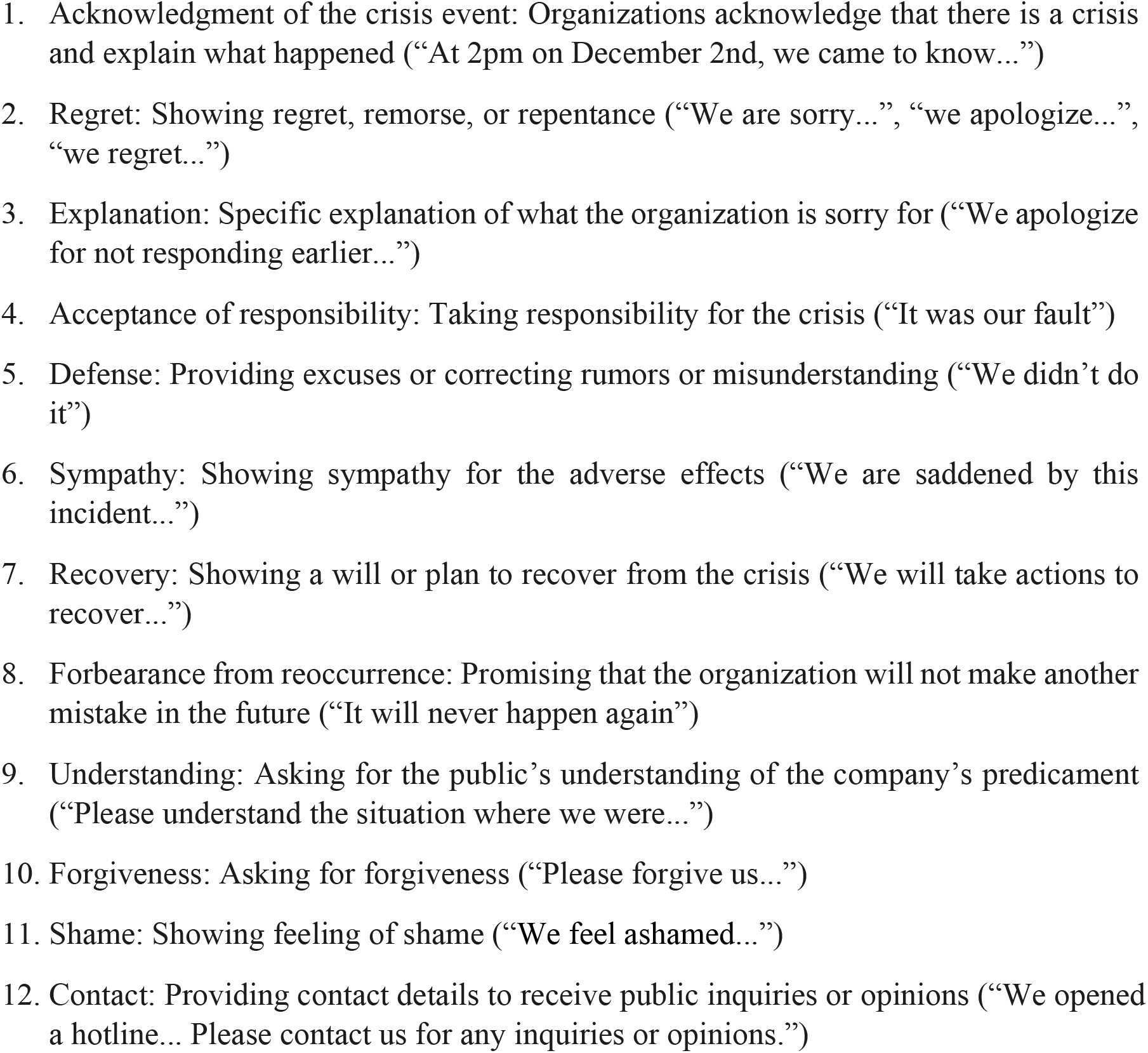
Content items found from our content analysis of actual public apologies issued.

The first eight elements match those from other content analyses of public apologies (Benoit, 1995; Boyd, 2011; Hearit, 2006; Page, 2014). In addition, we also found content elements 9-12. Whether they reflect cultural differences remains to be determined. For example, *shame* may be more culturally bound to South Korea, reflecting deep concern about social responsibility. Nonetheless, Munoz, chief executive of United Airlines, eventually said he felt “shame” when he saw the video of the violent removal of Dr. Dao from the plane (McCann, 2017). For our theoretical predictions, items 1-4 and 6-8 from the content analysis map directly to our predictions. Item 5, defense, relates generally to identifying causality and assigning responsibility, although more specifically relates to deflecting responsibility to others. Items 9-11 generally fall under a demonstration of true understanding via emotional statements (our prediction 2b), although in the content analysis they highlight the importance of a broad spectrum of related emotions to help influence the public’s evaluation of the company, event and its aftermath. Finally, item 12, contact, also may generally fall under redress of adverse consequences and forbearance; yet reveals a more specific example of how to apply it. The content analysis, then, provides real-world evidence for our theoretical predictions, and at the same time, offers additional specific content material to use in the apology statements.

In sum, theoretical and empirical work appear to point to greater effectiveness of more comprehensive public apologies, and content analyses of public apologies suggest that the derived content elements do make their way into various actual public apologies. However, whether comprehensive public apologies are indeed effective, and the actual content elements of the most effective ones — and thus recommendations for a public apology — remain unclear. We therefore sought to test our hypothesis that public apologies should generally be comprehensive, regardless of crisis severity, and also to determine exactly what an effective public apology should contain.

## Experiment 1: Public Apology Content

Because it is not reasonable to make a public apology based on single content elements, we began with a baseline “basic” public apology containing the minimal amount of necessary information. These elements were derived from our theoretical analysis as well as an attempt to be as neutral as possible, especially regarding responsibility, and thus liability, resulting in five elements: (1) what happened; (2) its significance; (3) why it happened; (4) how to prevent reoccurrence; (5) closing appeal for the public’s understanding (signifying respect and reconciliation). Basic apology examples (used for Experiments 1 & 2) are shown in Table 2.

**Table 2.**
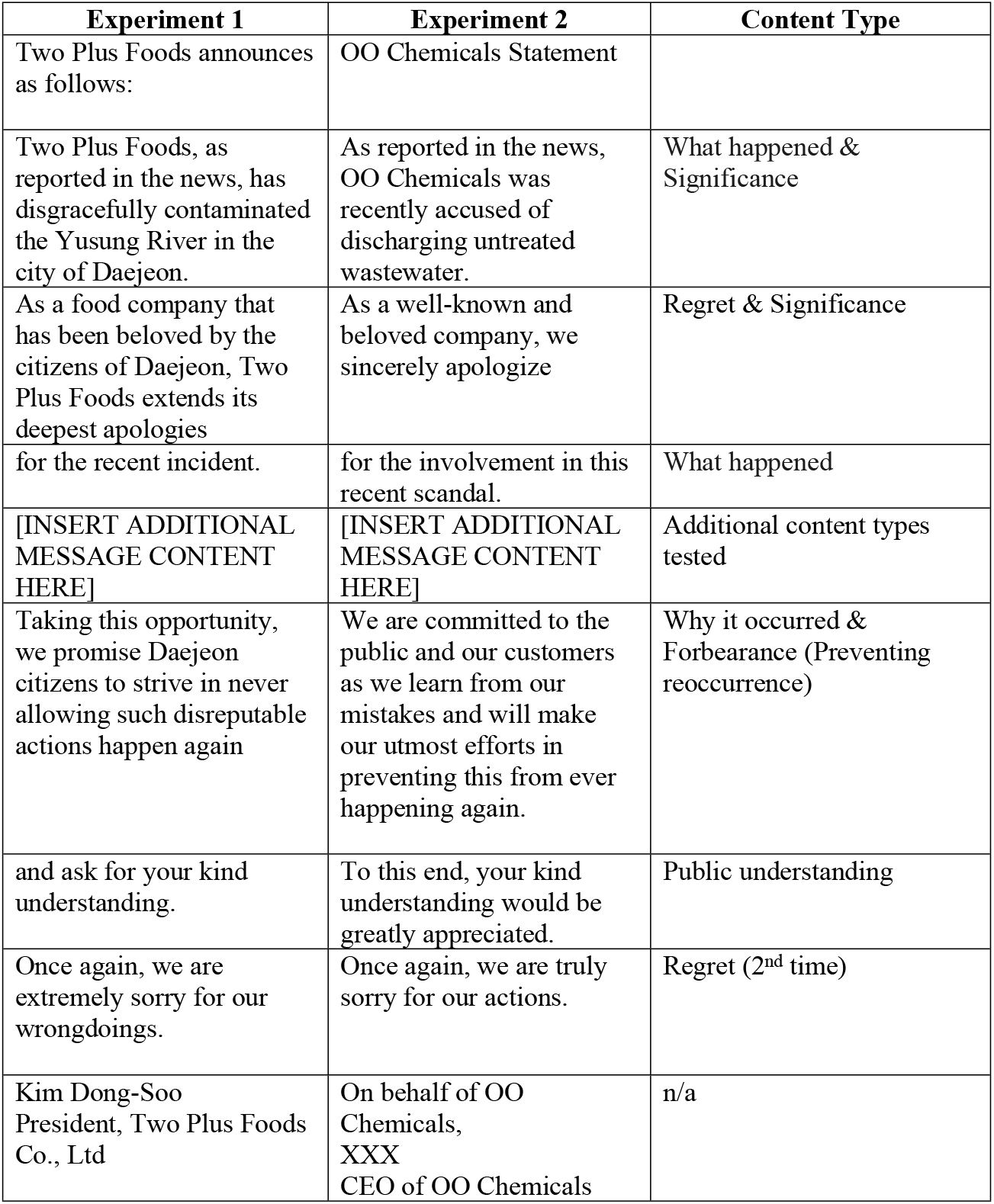
Basic apology statements used for Experiments 1 & 2 broken down by content type. The additional apology statements tested were created by inserting additional content elements in the fifth position.

We then tested the effectiveness of the basic apology statement as well as statements with additional content included. In Experiment 1 we tested all other content elements uncovered in our content analysis (list of 12) except for sympathy and forgiveness, which we considered similar enough to regret and understanding, respectively, to warrant setting aside for future study (Table 3) (Coombs & Holladay, 2008). To assess apology effectiveness, there are three main categories of responses that interact to determine outcome — cognition, affect, and behavior — and thus measures of each provide a more complete profile of effectiveness (Kim et al., 2015). More specifically, we used those listed in Table 3, which were adopted from (Coombs & Holladay, 2008).

**Table 3.**
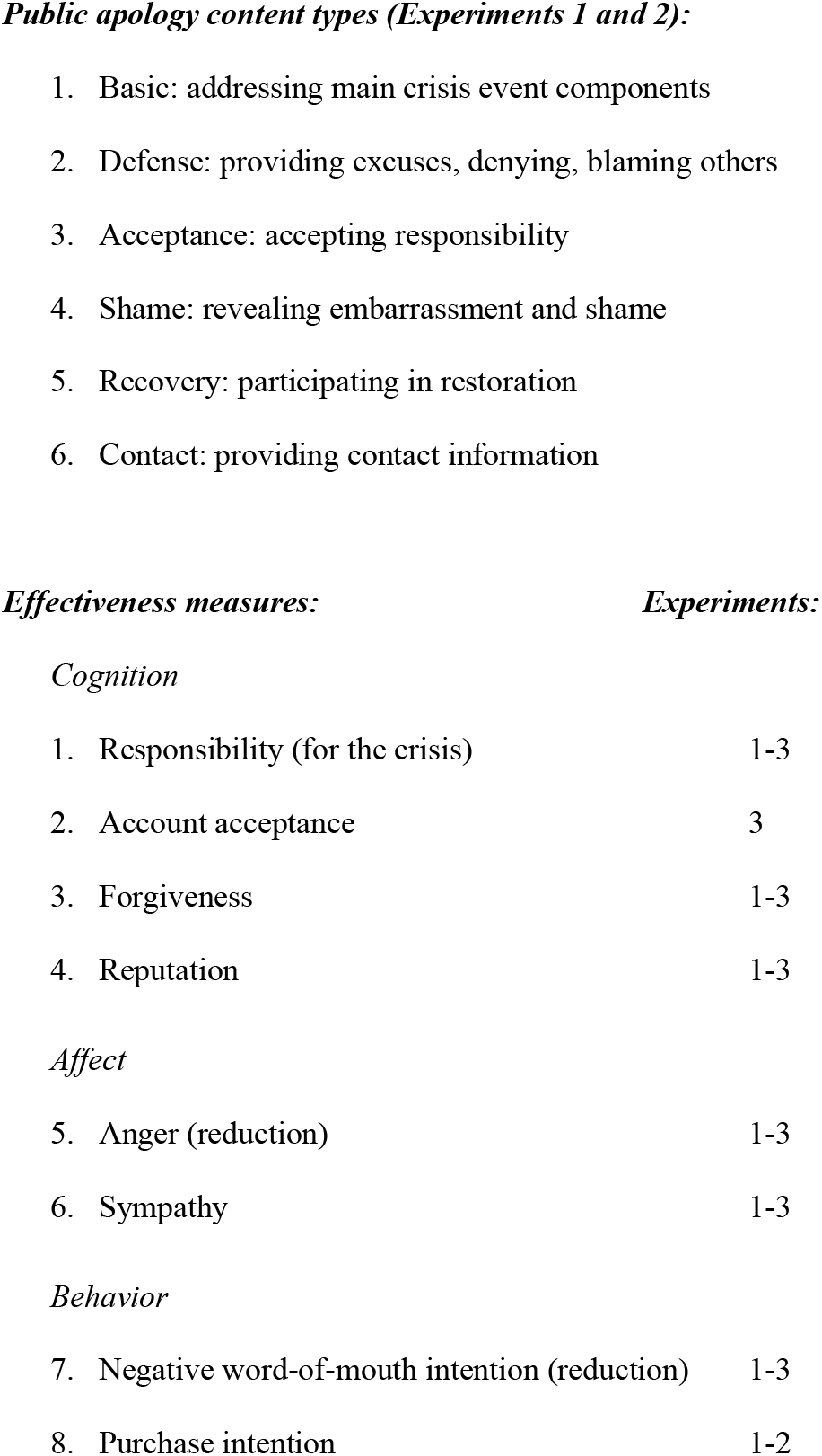
List of public apology content topics (independent variables) in Experiments 1 and 2, and participant responses (dependent measures of effectiveness, using a seven-point Likert-type scale) for all experiments.

|  |  |
| --- | --- |
| 1. Responsibility (for the crisis) | 1-3 |
| 2. Account acceptance | 3 |
| 3. Forgiveness | 1-3 |
| 4. Reputation | 1-3 |
| 5. Anger (reduction) | 1-3 |
| 6. Sympathy | 1-3 |
| 7. Negative word-of-mouth intention (reduction) | 1-3 |
| 8. Purchase intention | 1-2 |

Participants read a news report of a food processing company found to be polluting a regional river (see Table 1). As a control group, 300 college students participated in an initial online survey with no subsequent apology. They first read the news report and then were queried with seven effectiveness measures in Table 2 (all but account acceptance). For the main experimental manipulations, 230 college students participated in a laboratory study, where the participants read the same news report from a computer monitor, and then read one of six different public apology statements based on the content types listed in Table 2. We then queried the experimental group with the same seven effectiveness measures as the control group.

### Results

Table 4 shows the results with the experimental groups directly compared to the control group (i.e., no apology given). In sum, there were four key findings. First, the basic apology was effective, but only moderately so, being significant with all dependent measures together (“*All items*”), but when considering each effectiveness measure individually, only significantly improving *reputation* (thus 1 of 7). Second, on average (“*Experimental*” in Table 4), the enhanced apologies were effective, both for *All items* and for every individual dependent measure besides *responsibility*. Thus, in general, an enhanced message was superior to the basic one. Third, regarding the specific added statements, all five were effective when considering *All items* together. For individual message additions, (a) *acceptance* (of responsibility) improved all but one effectiveness measure (*responsibility*), (b) *defense, recovery*, and *contact* improved all but two (*responsibility* plus one other measure), and (c) *shame* improved only *reputation* and *anger (reduction)*. Thus, *acceptance* proved most effective, followed closely by *defense*, *recovery*, and *contact*. The acceptance content addition accepts full responsibility, including willingness to pay all liabilities. Given the high and similar degree of effectiveness for the four content topics vs. *shame*, we compared them to *shame*, and a commonality among the four appears to be having *actionable statements*, which demonstrate that measures are being taken to mitigate damages and/or correct the problem to minimize potential reoccurrence. The importance of taking actionable measures following the crisis further suggests that the adverse consequences (contaminating the water) and potential for reoccurrence are particularly important to the public (Bennett & Earwaker, 1994; Page, 2014; Scher & Darley, 1997; Schlenker & Darby, 1981). We therefore focused on these two factors in the subsequent experiments. Fourth, examining the individual dependent measures, *reputation* was the most changeable, improved by all apologies; whereas, *responsibility* could not be improved by any of the apologies. Thus, apologies are particularly effective in improving the public’s general perception of the wrongdoer (as measured by reputation); although exactly how this may influence future interactions is unclear, given that the other measures were less affected. Regarding crisis *responsibility*, clear statements about involvement and responsibility in the wrongdoing appear to solidify the public’s assessment of *responsibility* (even when the defense statement influences the other dependent measures — suggesting that participants still consider the organization to be ultimately responsibility).

**Table 4.**
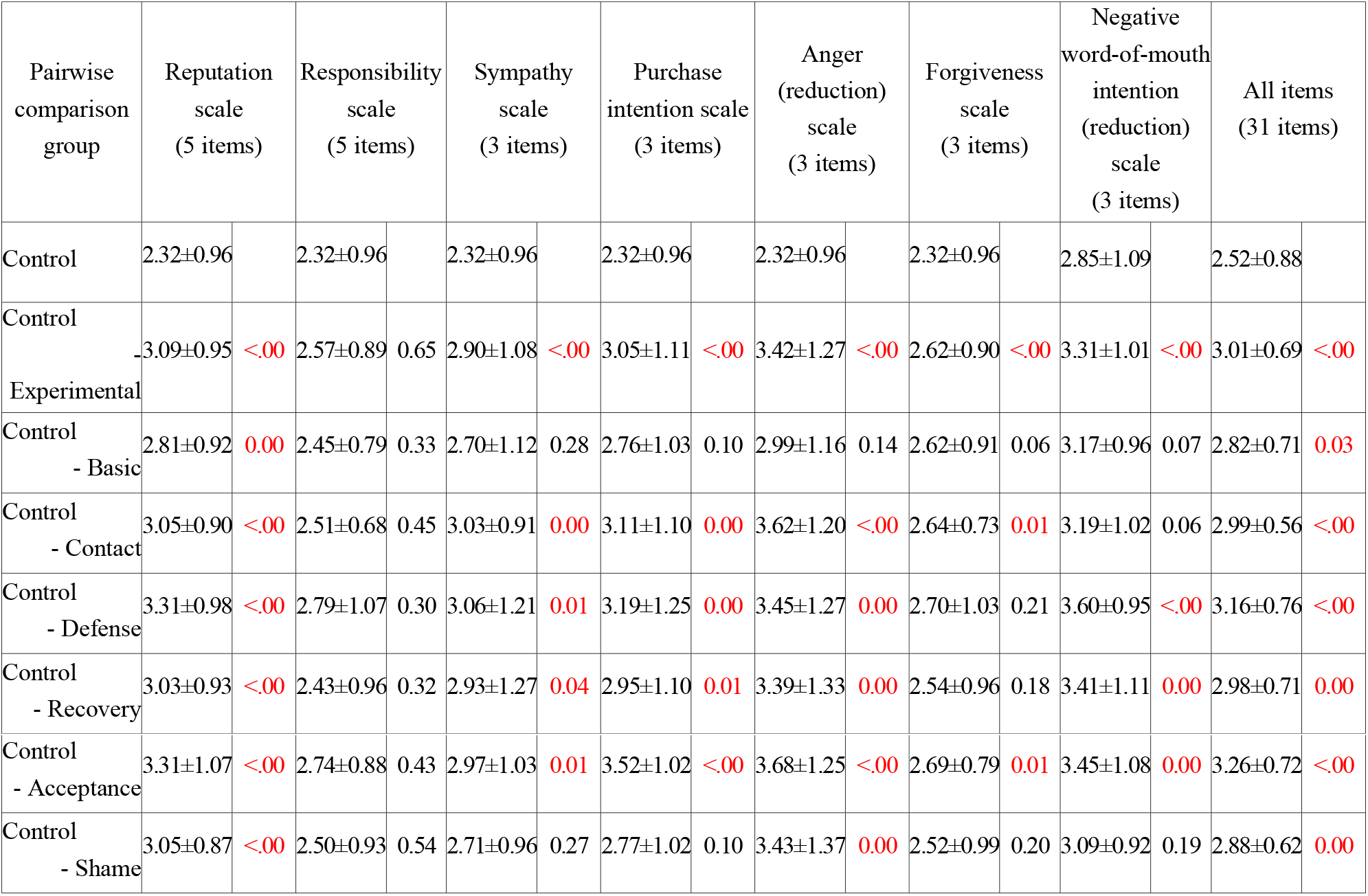
Experiment 1 crisis event results: river contamination by a food processing company (rating mean±std, Bonferroni corrected p-value from t-test comparison with no-apology control group; significant p-values in red).

## Experiment 2: More Harmful Crisis Event and Combinations of Content Additions to the Basic Apology

How the apology is received and therefore how it should be crafted may also depend on the severity of the crisis event (Bennett & Earwaker, 1994; Kim et al., 2004; Lewicki et al., 2016; Skarlicki et al., 2004; Struthers, Eaton, Santelli, Uchiyama & Shirvani, 2008); however, studies that have directly manipulated crisis severity are limited. In the interpersonal domain, two studies have found that as the consequence becomes more adverse (e.g., lightly bumping into someone vs. knocking them down and hurting them or accidental vs. irresponsible damaging of a watch), apology effectiveness decreases (Schlenker & Darby, 1981), with more elaborate apologies spontaneously given (Bennett & Earwaker, 1994). Nonetheless, the extent of the importance of crisis severity, especially in the corporate and public statement domain, remains largely unknown. Therefore, Experiment 2 tested whether consequence severity influences public apology effectiveness. Additionally, Experiment 2 examined public apology content further by testing the basic apology with the added statements from Experiment 1 as well as combinations of them.

In Experiment 1, we tested a basic apology statement as well as multiple enhanced versions and found that not only did additional messaging improve effectiveness, but that two factors of the crisis event itself appear to be particularly important to people: the adverse effects resulting from the crisis and potential for reoccurrence. In Experiment 2, we focused on *adverse consequences* and attempted to use a more harmful situation, with the headline reading “00 Chemicals accused of discharging carcinogenic and toxic substances” (versus general waste contamination by a food company in Experiment 1). As discussed, prior studies have found evidence for an inverse relationship between crisis severity and apology effectiveness during interpersonal conflict (Bennett & Earwaker, 1994; Schlenker & Darby, 1981), however, its significance with respect to corporate crises remains unclear. Moreover, in addition to testing apology effectiveness as done in Experiment 1, we also tested combinations of the five content types, for a total of 22 different content additions to the basic apology. The experiment was conducted as an online survey of 2,800 participants, of which 600 were used as a control group, being exposed to the news report without any apology. All participants were queried using the same seven dependent measures used in Experiment 1 (Table 3).

### Results

Table 5 shows the results. In sum, there again were four main findings. First, the basic apology was not effective when considering all dependent measures together (*All items*), and was only significant for one measure separately (*purchase intent*). Second, as was found in Experiment 1, on average, the enhanced apologies were effective, both for *All items* and for every individual dependent measure besides *responsibility*. Thus, an enhanced message was again superior to the basic one. Third, with respect to the specific message additions — i.e., each experimental condition separately — one condition stood out as particularly effective: the apology with the added combination of *recovery* and *defense* statements. When these two additions were made, every effectiveness measure was improved except for *responsibility* (thus, 6 of 7 individual measures; with the next highest being 3 of 7, achieved by three other messages). Fourth, regarding the individual dependent measures, as in Experiment 1, *reputation* was the most pliable, improved by multiple content additions, whereas *responsibility* was not affected by any public apology statements.

**Table 5.**
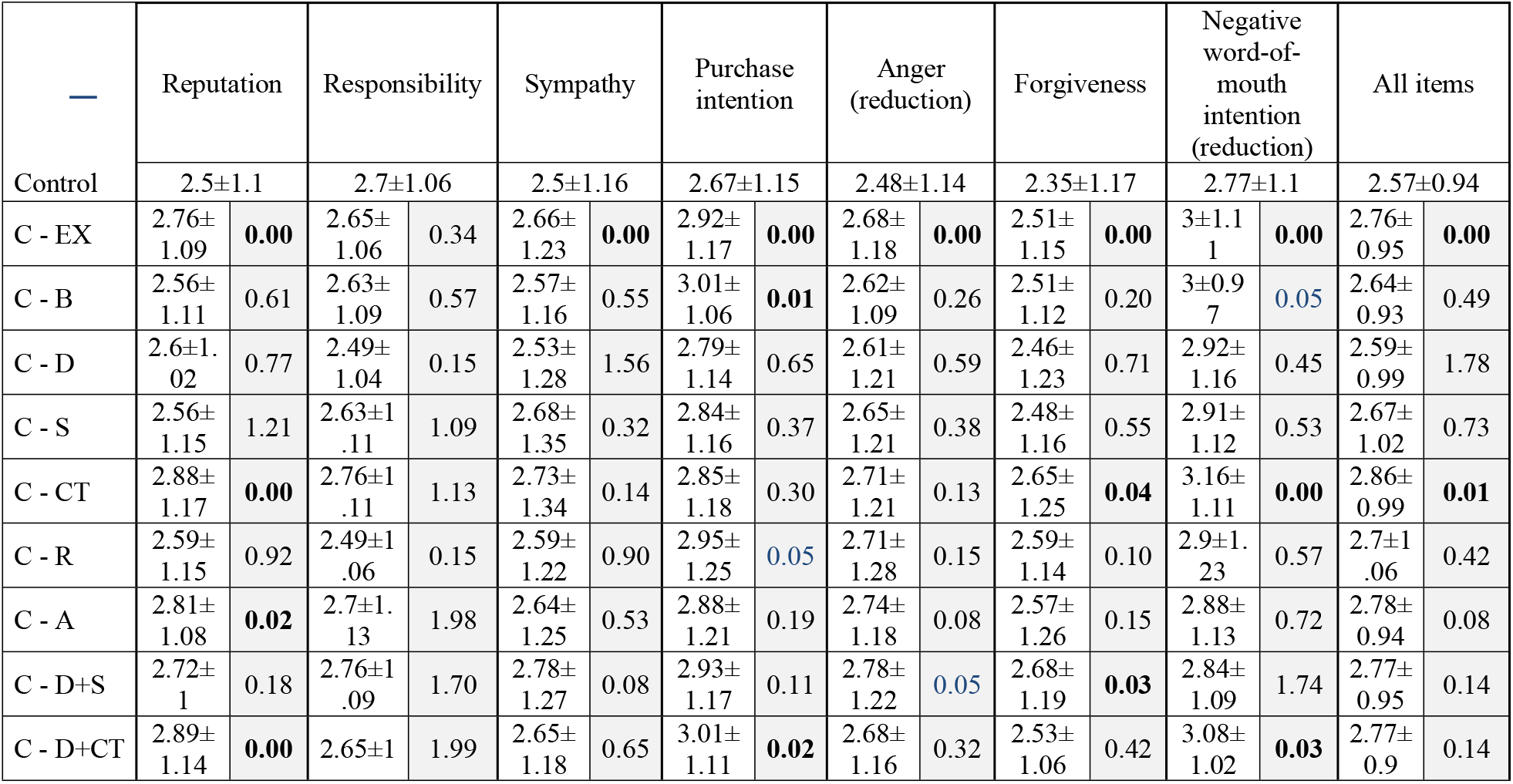
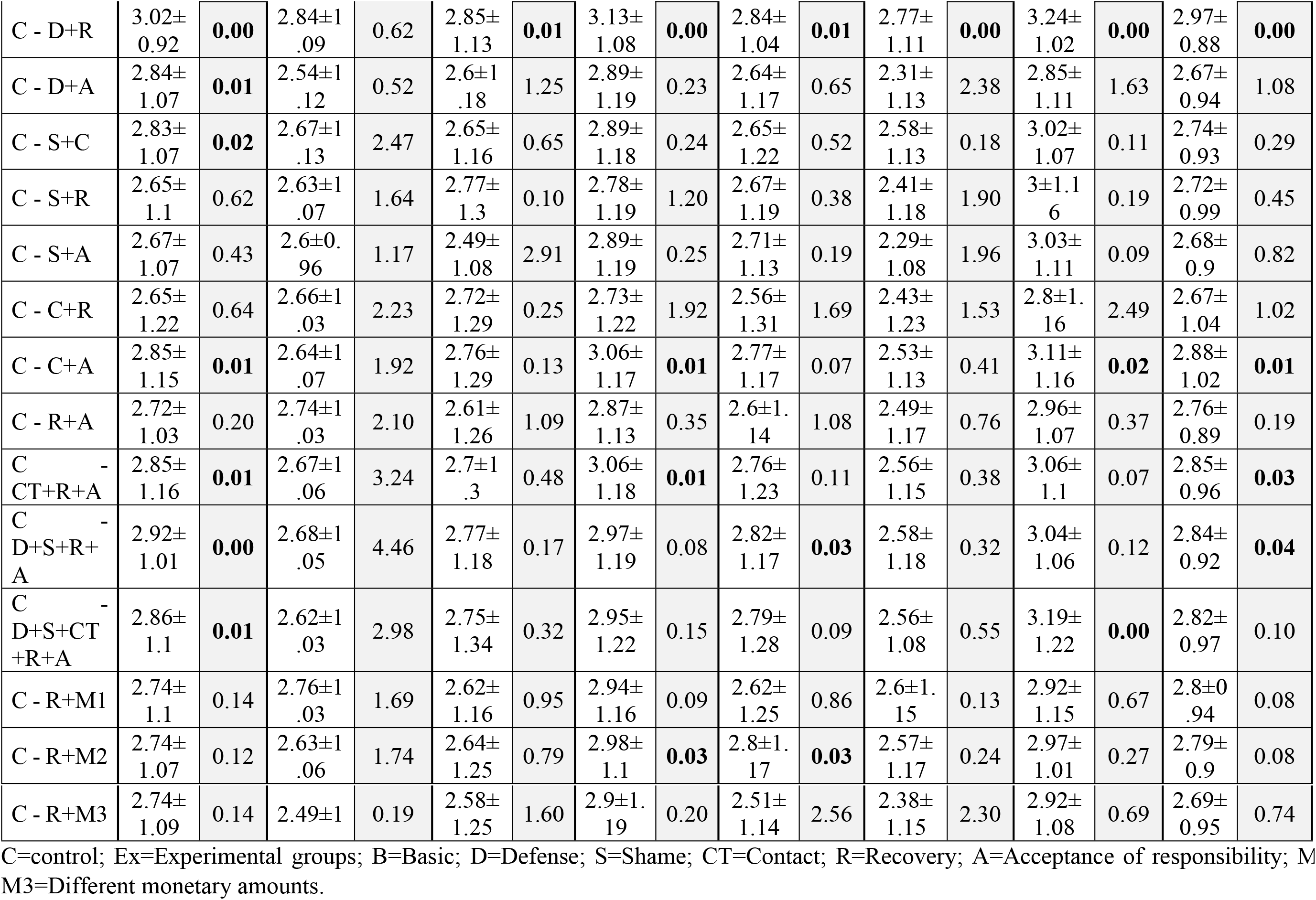
Experiment 2 crisis event results: river contamination by a chemical company (rating mean±std, Bonferroni corrected p-value from t-test comparison with no-apology control group; significant p-values in bold).

| — | Reputation |  | Responsibility |  | Sympathy |  | Purchase intention |  | Anger (reduction) |  | Forgiveness |  | Negative word-of-mouth intention (reduction) |  | All items |  |
| --- | --- | --- | --- | --- | --- | --- | --- | --- | --- | --- | --- | --- | --- | --- | --- | --- |
| Control | 2.5±1.1 | | 2.7±1.06 | | 2.5±1.16 | | 2.67±1.15 | | 2.48±1.14 | | 2.35±1.17 | | 2.77±1.1 | | 2.57±0.94 | |
| C - EX | 2.76±<br>1.09 | **0.00** | 2.65±<br>1.06 | 0.34 | 2.66±<br>1.23 | **0.00** | 2.92±<br>1.17 | **0.00** | 2.68±<br>1.18 | **0.00** | 2.51±<br>1.15 | **0.00** | 3±1.1<br>1 | **0.00** | 2.76±<br>0.95 | **0.00** |
| C - B | 2.56±<br>1.11 | 0.61 | 2.63±<br>1.09 | 0.57 | 2.57±<br>1.16 | 0.55 | 3.01±<br>1.06 | **0.01** | 2.62±<br>1.09 | 0.26 | 2.51±<br>1.12 | 0.20 | 3±0.9<br>7 | **0.05** | 2.64±<br>0.93 | 0.49 |
| C - D | 2.6±1.<br>02 | 0.77 | 2.49±<br>1.04 | 0.15 | 2.53±<br>1.28 | 1.56 | 2.79±<br>1.14 | 0.65 | 2.61±<br>1.21 | 0.59 | 2.46±<br>1.23 | 0.71 | 2.92±<br>1.16 | 0.45 | 2.59±<br>0.99 | 1.78 |
| C - S | 2.56±<br>1.15 | 1.21 | 2.63±1<br>.11 | 1.09 | 2.68±<br>1.35 | 0.32 | 2.84±<br>1.16 | 0.37 | 2.65±<br>1.21 | 0.38 | 2.48±<br>1.16 | 0.55 | 2.91±<br>1.12 | 0.53 | 2.67±<br>1.02 | 0.73 |
| C - CT | 2.88±<br>1.17 | **0.00** | 2.76±1<br>.11 | 1.13 | 2.73±<br>1.34 | 0.14 | 2.85±<br>1.18 | 0.30 | 2.71±<br>1.21 | 0.13 | 2.65±<br>1.25 | **0.04** | 3.16±<br>1.11 | **0.00** | 2.86±<br>0.99 | **0.01** |
| C - R | 2.59±<br>1.15 | 0.92 | 2.49±1<br>.06 | 0.15 | 2.59±<br>1.22 | 0.90 | 2.95±<br>1.25 | **0.05** | 2.71±<br>1.28 | 0.15 | 2.59±<br>1.14 | 0.10 | 2.9±1.<br>23 | 0.57 | 2.7±1<br>.06 | 0.42 |
| C - A | 2.81±<br>1.08 | **0.02** | 2.7±1.<br>13 | 1.98 | 2.64±<br>1.25 | 0.53 | 2.88±<br>1.21 | 0.19 | 2.74±<br>1.18 | 0.08 | 2.57±<br>1.26 | 0.15 | 2.88±<br>1.13 | 0.72 | 2.78±<br>0.94 | 0.08 |
| C - D+S | 2.72±<br>1 | 0.18 | 2.76±1<br>.09 | 1.70 | 2.78±<br>1.27 | 0.08 | 2.93±<br>1.17 | 0.11 | 2.78±<br>1.22 | **0.05** | 2.68±<br>1.19 | **0.03** | 2.84±<br>1.09 | 1.74 | 2.77±<br>0.95 | 0.14 |
| C - D+CT | 2.89±<br>1.14 | **0.00** | 2.65±1 | 1.99 | 2.65±<br>1.18 | 0.65 | 3.01±<br>1.11 | **0.02** | 2.68±<br>1.16 | 0.32 | 2.53±<br>1.06 | 0.42 | 3.08±<br>1.02 | **0.03** | 2.77±<br>0.9 | 0.14 |
| C - D+R | 3.02±<br>0.92 | **0.00** | 2.84±1<br>.09 | 0.62 | 2.85±<br>1.13 | **0.01** | 3.13±<br>1.08 | **0.00** | 2.84±<br>1.04 | **0.01** | 2.77±<br>1.11 | **0.00** | 3.24±<br>1.02 | **0.00** | 2.97±<br>0.88 | **0.00** |
| C - D+A | 2.84±<br>1.07 | **0.01** | 2.54±1<br>.12 | 0.52 | 2.6±1<br>.18 | 1.25 | 2.89±<br>1.19 | 0.23 | 2.64±<br>1.17 | 0.65 | 2.31±<br>1.13 | 2.38 | 2.85±<br>1.11 | 1.63 | 2.67±<br>0.94 | 1.08 |
| C - S+C | 2.83±<br>1.07 | **0.02** | 2.67±1<br>.13 | 2.47 | 2.65±<br>1.16 | 0.65 | 2.89±<br>1.18 | 0.24 | 2.65±<br>1.22 | 0.52 | 2.58±<br>1.13 | 0.18 | 3.02±<br>1.07 | 0.11 | 2.74±<br>0.93 | 0.29 |
| C - S+R | 2.65±<br>1.1 | 0.62 | 2.63±1<br>.07 | 1.64 | 2.77±<br>1.3 | 0.10 | 2.78±<br>1.19 | 1.20 | 2.67±<br>1.19 | 0.38 | 2.41±<br>1.18 | 1.90 | 3±1.1<br>6 | 0.19 | 2.72±<br>0.99 | 0.45 |
| C - S+A | 2.67±<br>1.07 | 0.43 | 2.6±0.<br>96 | 1.17 | 2.49±<br>1.08 | 2.91 | 2.89±<br>1.19 | 0.25 | 2.71±<br>1.13 | 0.19 | 2.29±<br>1.08 | 1.96 | 3.03±<br>1.11 | 0.09 | 2.68±<br>0.9 | 0.82 |
| C - C+R | 2.65±<br>1.22 | 0.64 | 2.66±1<br>.03 | 2.23 | 2.72±<br>1.29 | 0.25 | 2.73±<br>1.22 | 1.92 | 2.56±<br>1.31 | 1.69 | 2.43±<br>1.23 | 1.53 | 2.8±1.<br>16 | 2.49 | 2.67±<br>1.04 | 1.02 |
| C - C+A | 2.85±<br>1.15 | **0.01** | 2.64±1<br>.07 | 1.92 | 2.76±<br>1.29 | 0.13 | 3.06±<br>1.17 | **0.01** | 2.77±<br>1.17 | 0.07 | 2.53±<br>1.13 | 0.41 | 3.11±<br>1.16 | **0.02** | 2.88±<br>1.02 | **0.01** |
| C - R+A | 2.72±<br>1.03 | 0.20 | 2.74±1<br>.03 | 2.10 | 2.61±<br>1.26 | 1.09 | 2.87±<br>1.13 | 0.35 | 2.6±1.<br>14 | 1.08 | 2.49±<br>1.17 | 0.76 | 2.96±<br>1.07 | 0.37 | 2.76±<br>0.89 | 0.19 |
| C<br>CT+R+A | 2.85±<br>1.16 | **0.01** | 2.67±1<br>.06 | 3.24 | 2.7±1<br>.3 | 0.48 | 3.06±<br>1.18 | **0.01** | 2.76±<br>1.23 | 0.11 | 2.56±<br>1.15 | 0.38 | 3.06±<br>1.1 | 0.07 | 2.85±<br>0.96 | **0.03** |
| C<br>D+S+R+<br>A | 2.92±<br>1.01 | **0.00** | 2.68±1<br>.05 | 4.46 | 2.77±<br>1.18 | 0.17 | 2.97±<br>1.19 | 0.08 | 2.82±<br>1.17 | **0.03** | 2.58±<br>1.18 | 0.32 | 3.04±<br>1.06 | 0.12 | 2.84±<br>0.92 | **0.04** |
| C<br>D+S+CT<br>+R+A | 2.86±<br>1.1 | **0.01** | 2.62±1<br>.03 | 2.98 | 2.75±<br>1.34 | 0.32 | 2.95±<br>1.22 | 0.15 | 2.79±<br>1.28 | 0.09 | 2.56±<br>1.08 | 0.55 | 3.19±<br>1.22 | **0.00** | 2.82±<br>0.97 | 0.10 |
| C - R+M1 | 2.74±<br>1.1 | 0.14 | 2.76±1<br>.03 | 1.69 | 2.62±<br>1.16 | 0.95 | 2.94±<br>1.16 | 0.09 | 2.62±<br>1.25 | 0.86 | 2.6±1.<br>15 | 0.13 | 2.92±<br>1.15 | 0.67 | 2.8±0<br>.94 | 0.08 |
| C - R+M2 | 2.74±<br>1.07 | 0.12 | 2.63±1<br>.06 | 1.74 | 2.64±<br>1.25 | 0.79 | 2.98±<br>1.1 | **0.03** | 2.8±1.<br>17 | **0.03** | 2.57±<br>1.17 | 0.24 | 2.97±<br>1.01 | 0.27 | 2.79±<br>0.9 | 0.08 |
| C - R+M3 | 2.74±<br>1.09 | 0.14 | 2.49±1 | 0.19 | 2.58±<br>1.25 | 1.60 | 2.9±1.<br>19 | 0.20 | 2.51±<br>1.14 | 2.56 | 2.38±<br>1.15 | 2.30 | 2.92±<br>1.08 | 0.69 | 2.69±<br>0.95 | 0.74 |

C=control; Ex=Experimental groups; B=Basic; D=Defense; S=Shame; CT=Contact; R=Recovery; A=Acceptance of responsibility; M1-M3=Different monetary amounts.

To assess crisis severity, we compare Experiments 1 and 2. For the basic apology statement, its moderate effectiveness in Experiment 1 (for *All items* and *reputation*) reduced to even weaker effectiveness in Experiment 2 (significant only for *purchase intent)*. Moreover, although in both experiments, additional content elements generally improved effectiveness, the most effective addition differed in the two experiments: in the less severe case of Experiment 1, accepting *responsibility* was most effective, with three others comparable as well; whereas for the more severe case in Experiment 2, *defense* and *recovery* combined was by far the most effective addition. Thus, crisis severity had a major impact on how the apologies were received. In fact, in a broader context, even river contamination by a food processing company (Experiment 1) might be considered intermediately severe, given that the basic apology was only moderately effective, requiring enhancement by additional statements. Thus, for an even less severe crisis, the basic apology might be expected to show greater effectiveness. In any case, it is clear that crisis severity significantly impacts apology effectiveness and therefore how apologies should be constructed. Our findings thus extend those that have found such a relationship in interpersonal cases to corporate crises and public apologies made following more impactful transgressions (Bennett & Earwaker, 1994; Schlenker & Darby, 1981).

Severity appears to influence the degree of public concern, and thus the extent to which they engage in the specifics of the crisis and its aftermath. When in less severe cases, additional *actionable* statements may induce a sense that the wrongdoer is taking sufficient steps to make amends and limit potential reoccurrence. In more severe cases, the target audience appears less willing to accept a basic account, and further, appears to require more specific statements — in this case, regarding both reparations and defense. It certainly seems reasonable that when a clear and present danger exists, such as carcinogens in the drinking water, concrete steps to eliminate it (i.e., cleanup and recovery) would be important (Coombs & Holladay, 2008; Lewicki et al., 2016; Page, 2014; Scher & Darley, 1997).

The reason for the defense message effectiveness, however, is less clear, given that the statement both deflected responsibility (“…we have not been regularly polluting but rather *Water Cleaner Inc.*, our vendor responsible for the treatment of our sewage, is at fault for the latest discovery.”) and took measures to prevent the actual wrongdoer from repeating the offense (“…we plan on reinforcing supervision of our vendor moving forward.”). Thus, it both pointed to an external cause but also suggested control over it, to eliminate future reoccurrence. It therefore is unclear which factor may have been particularly effective. The next experiment thus tests these factors (external source versus the ability to control it) more directly.

## Experiment 3: Control Over the Crisis Source

Results from Experiments 1 (multiple effective statements pointing to actions taken) and 2 (recovery and defense) suggest that people may be especially concerned about future threats and the actions taken to mitigate them. If so, it may imply that we are sensitive to the specific causes of the crisis and whether or not they can be controlled by the organization. Two factors are particularly relevant for assessing long-term causal control of the crisis source: (1) whether the company can indeed potentially control the source (i.e., there is a clear causal link between them), i.e., *controllability*; and (2) the presence of other factors that could also influence the source and thus reduce or negate the company’s potential influence. Regarding factor (2), whether the crisis source is *internal* or *external* to the company is particularly relevant given that an internal source should be less susceptible to other possible external influences as well as more readily monitored and controlled when it is within the organization’s normal purview. (From a Bayesian inference perspective, factor one reflects the *likelihood* and factor two reflects the *prior* relative to the *marginal*.) Thus, having the source of the crisis as both *internal* to the organization and *controllable* would provide the strongest case for organizational control over the causal factors.

Therefore, we tested an *“internal and controllable (IC)* causal attributions” condition versus an *“external and uncontrollable (EU)”* condition on public perceptions of organizations involved in crises. By *causal attributions* we mean assessment of both (a) the cause of the crisis (internal vs. external) and (b) controllability over the causal source by the organization (Lee et al., 2004). With respect to the cause being internal vs. external to the organization, we focused on the causal *actions* leading to the wrongdoing and whether they were part of the normal operating procedures of the company. Thus, the wrongdoer could be an employee of the organization, but whether the wrongdoing was considered internal or external depended on the particular actions taken. Furthermore, we recognized two types of causal actions that should both be considered internal to, and thus the direct result of, the organization: (1) actions taken as part of the standard operating practices of the organization that directly produced the harmful event; or (2) actions *not* taken but should have been taken as part of standard operating practices, which would have prevented the harmful event. Thus, case (1) or (2) were considered internal to the organization, otherwise they were classified as external. Table 6 provides an example of case (1) for the internal attribution.

**Table 6.** Sample crisis incident and public apologies used for the internal-controllable and external-uncontrollable attribution conditions. Faulty ignition leads to a flood of complaints against OO Motors OO Motor’s latest model launched earlier this year was found to have problems with ignition caused by defects in its electric cabling. However, OO Motors has been keeping quiet about the ignition failures while only fixing it for customers who had complained leading to a potential class action started online. Meanwhile, OO Motors rebutted the fact claiming it was just a common problem. Consumer complaints are unlikely to dissipate for a while. (Industrial News, Kim Hyung-Kyun)

|  | Regret | External / Internal | Uncontrollable / Controllable | Repetition & Closing |
| --- | --- | --- | --- | --- |
| External/<br>Uncontrollable | Apology from OO Motors:<br>We are truly sorry for the recent controversy spread across the web and in the media regarding our latest model OO. | An internal investigation by OO Motors has confirmed that the problem was caused by quality issues at our outsource partner that has been supplying parts for several of the 2010 models.<br>**EXTERNAL:**<br>**Blaming**<br>**“outsource partner”** | The failure was a result of a part manufactured by an outsource partner therefore it was hard for us to take preventive measures. | OO Motors is committed to always strive for customer safety.<br>OOO<br>CEO of OO Motors |
| Internal/<br>Controllable |  | An internal investigation by OO Motors has proven that there were problems with some of the parts used in the 2010 models. | As a company that should have been responsible for final quality assurance we have failed to thoroughly control the process. |  |
|  |  | INTERNAL: No<br>suggestion that<br>the cause was<br>external |  |  |

### Results

As seen in Table 7, the apology effectiveness ratings were (a) higher for the *internal-controllable* attributions condition than for the *external-uncontrollable* condition for all seven dependent measures (nonparametric sign test, p=0.016); (b) significantly higher *overall*, as well as for *account acceptance*, *sympathy*, *anger*, and *reputation*; and (c) approaching significance for *negative word-of-mouth intention reduction* (p=0.055). Thus, when the source of the crisis was more clearly under the control of the wrongdoer, the public appears to be more convinced by the apology. This result suggests that the potential threat of crisis reoccurrence is particularly important, with the public finely tuned to the actual causal factors of the crisis event and whether they can be controlled by the organization.

**Table 7.** Comparison of the rating means between external/uncontrollable (EU) and internal/controllable (IC) public apologies on a seven-point scale (*: p<0.05; **: p<0.01; ***: p<0.001).

|  | *Overall* |  | *Account acceptance*<br>The people affected by the incident would consider the response by the organization to be appropriate. |  | *Responsibility (for the crisis)*<br>The blame for the incident lies with the organization. |  | *Sympathy*<br>Because of the incident situation, I feel sorry for the organization. |  | *Forgiveness*<br>The organization should be punished for the incident. (Inverted scale) |  | *Negative word-of-mouth intention (reduction)*<br>I would encourage friends or relatives not to buy products or services from the organization. |  | *Anger (reduction)*<br>I feel annoyed toward the organization for what happened. |  | *Reputation*<br>In most situations, I would trust what the organization said. |  |
| --- | --- | --- | --- | --- | --- | --- | --- | --- | --- | --- | --- | --- | --- | --- | --- | --- |
|  | EU | IC | EU | IC | EU | IC | EU | IC | EU | IC | EU | IC | EU | IC | EU | IC |
| Mean | 2.66 | 2.89 | 2.52 | 3.16 | 2.48 | 2.50 | 2.48 | 2.79 | 2.57 | 2.63 | 3.28 | 3.41 | 2.65 | 2.83 | 2.62 | 2.93 |
| p-value | 0.00*** |  | 0.00*** |  | 0.71 |  | 0.00*** |  | 0.39 |  | 0.06 |  | 0.01** |  | 0.00*** |  |
| STD | 1.13 | 1.23 | 1.48 | 1.74 | 1.49 | 1.39 | 1.45 | 1.58 | 1.48 | 1.46 | 1.61 | 1.70 | 1.45 | 1.53 | 1.40 | 1.51 |

Our results support those found for annual reports to stockholders who must determine whether to continue investing in a company based on the reports (Lee et al., 2004). When negative events are explained as being based on causes firmly within the company’s control (i.e., both internal and controllable), faith remains higher than if out-of-control, even if that means the company made mistakes that led to adverse consequences. This suggests that future performance is most critical to stockholders, with controllability strongly implying overall competence, with the expectation that it will lead to fewer problems and better performance in the future. Our results thus extend the findings found for annual reports and future stock performance to crisis events and public apologies in general, and how cognition, affect, and behavior of the general public are influenced by causal attributions.

Given that the causal information was communicated by the organization itself rather than via an outside source, the identification of an internal source may also promote account believability and organizational integrity (by implicating oneself). However, given (a) the success of the defense statements in both Experiments 1 and 2, in which an external source is implicated but with control over it, and (b) the greater effectiveness here of a controllable, internal source (IC) over an uncontrollable, external source (EU), controllability appears to be a dominant factor (see Lee et al., 2004). Thus, taken together, our results point to the effectiveness of delineating the causal factors and their controllability in the apology. This effectiveness in turn underscores the importance of threats and the likelihood of crisis reoccurrence to the public. Future studies should therefore examine the role of causal attributions more thoroughly, such as comparing the IC and EU conditions when the causal details are reported by an external source (e.g., media) versus by the organization itself, and whether some cases may still benefit from responsibility deflection (such as with extremely adverse consequences): that is, the potential effectiveness of implicating external sources that are yet controllable by the organization.

## General Discussion

In both Experiments 1 and 2 there was some evidence for the effectiveness of the basic apology, but it was readily improved with additional content. This improvement suggests that increasingly consequential crises may lead to greater engagement by the audience in the details of the crisis, requiring increasingly thoughtful apologies (beyond the basic apology) (Schlenker & Darby, 1981). In Experiment 1 the common theme among the most effective additions appeared to involve *actual actions taken* by the wrongdoer to help rectify matters, which further implies that people may be particularly sensitive to mitigating the harm induced (such as via recovery efforts and reparations), and more generally to rectifying the fairness imbalance, and removing the potential lingering threat (potential reoccurrence).

Given that crisis severity likely influences how people react to crises, Experiment 2 examined its significance by increasing the hazard of the river contaminant — via potentially carcinogenic chemicals. The experiment also tested multiple combinations of additional apology messaging. One apology statement stood out as superior: the basic apology plus recovery and defense. The difference in the effective message additions between Experiments 1 and 2 shows that crisis severity impacts apology effectiveness. The findings thus extend others that found an effect of crisis severity with interpersonal crises (bumping into people, damaging someone’s property) to corporate crises and public apologies (Bennett & Earwaker, 1994; Schlenker & Darby, 1981). Thus, there is no ‘one size fits all’ apology, and crisis specifics must be taken into account when formulating the message (Bennett & Earwaker, 1994; Schlenker & Darby, 1981; Schumann, 2014). In the more severe crisis of Experiment 2, a statement regarding specific actions taken to reverse the resulting damages was needed, together with one on defense. In cases with looming danger, such as chemical contamination of the local water, concrete measures taken to remove the hazard (e.g., via restoration) would certainly be expected to be effective (Lewicki & Polin, 2012; Lewicki et al., 2016; Minow, 2009; Schumann, 2014). With respect to a strategy of defense, both Experiments 1 and 2 found evidence for its effectiveness. Thus, our results may support and extend others that suggest defense may be an effective strategy under more egregious cases, such as when the violation was intended (Kim et al., 2004) or when consequences are especially onerous. However, in our case, exactly what aspect of the defense statement most underpinned its effectiveness remains unclear: i.e., whether deflection of responsibility or action taken to prevent crisis reoccurrence.

Because our defense statements (in both Experiments 1 and 2) contained a message of controllability over the cause of the crisis, and because the other findings (actions taken in Experiment 1, recovery in Experiment 2) point to the importance of threat reduction to the public, we examined this concern in Experiment 3. Statements clarifying causality (i.e., specific actions taken or inactions that should have been taken that led to the wrongdoing) may help convince people that the source of the crisis is properly identified and the problem resolved; and Experiment 3 tested this hypothesis and indeed found evidence for its importance in apology assessment (Lee et al., 2004). In fact, when considering the primary concerns people have in the face of crises (e.g., why did it occur, who was responsible, what were the adverse consequences, how will the harm be mitigated, what level of retribution is appropriate for the wrongdoing with respect to fairness and social justice, what is the likelihood of reoccurrence), all major concerns appear to require a clear understanding of the crisis event details for proper assessment and subsequent apology effectiveness. Thus, people appear most convinced when they can assess for themselves the event details and subsequent measures taken. This finding supports the view that apologies may generally be met with suspicion unless material evidence is provided (Coombs & Holladay, 2008; De Cremer, 2010; Farrell & Rabin, 1996), and further suggests that causal attributions play a fundamental role in people’s ultimate assessment of the crisis event and its aftermath.

Taken together, our evidence suggests that an optimal public statement reflects the enormity of the crisis, but nevertheless is comprehensive, attempting to address all of the public’s concerns (Lewicki et al., 2016; Schumann, 2014). It does so by expressing a sincere conviction to rectify matters (Schumann, 2014; Tomlinson et al., 2004; Zechmeister et al., 2004), and an ability to do so via (a) a specification of the causal factors and how they are controllable, and (b) statements of actual actions taken to prevent reoccurrence and to repay the debt owed to the victims (Kim et al., 2015).

‘Sorry’ — even for companies in crisis — is indeed not nearly enough.

## Methods

### Experiment 1

#### Participants

530 college students took part in this study: 47% female (n=108) and 53% male (n=122), with average age 24.28. All subjects were recruited from Daejeon, South Korea and were paid for participating (KRW 12,000 — approximately $10.70). 300 control subjects participated only in the online behavioral survey, which was conducted by panel members of a professional research firm in Seoul, Korea. The survey participants earned cash points. The participants were 54.3% female (n=163) and 45.7% male (n=137), with average age 22.3 (median=22.0). The study was carried out in accordance with the recommendations of the Institutional Review Board of KAIST, and the protocol was approved by them.

#### Materials

A mock news story was developed for the experiment by one of the authors and then revised by a professional journalist in South Korea. The news story was about a fake food company called Two Plus Food, which had repeatedly polluted Yuseong cheon, a major stream in Daejeon, for the last two years. From Coombs (2007), crises can be organized into three categories: “victim crisis”, when companies have minimal crisis responsibility, such as with natural disasters or product tampering; “accidental crisis”, when companies have low crisis responsibility, such as with technical error accidents or technical error product harm; and “preventable crisis”, when companies have strong crisis responsibility, such as with organizational misdeeds (e.g., intentional environment pollution or human-error product harm). For a strong test of apology effectiveness, the scenario used in this study was a *preventable crisis*, where the company appeared to have strong responsibility for the crisis.

The experimental group was divided into six subgroups in which the first received the basic public apology (Table 1) and the other five an apology with an additional content element: basic + defense (providing excuses for wrongdoing); basic + recovery (“we are doing…to recover…”); basic + acceptance (of responsibility) (“we are responsible…”); basic + shame (“we feel ashamed…”); basic + recovery (“we are doing…to recover…”); basic + contact (“please contact… for any inquiries”);. The added message components were placed in the same location, between the second and the third paragraphs, of each basic apology statement.

All responses were measured using the questionnaire from Coombs and Holladay (2008) developed by them and others (Blumstein et al., 1974; Coombs & Holladay, 2002; Griffin, Babin & Darden, 1992; Jorgensen, 1996; McAuley, Duncan & Russell, 1992), translated into Korean. As shown in Table 2, we measured seven dimensions: three cognitive, crisis responsibility, forgiveness, and company reputation; two affective, anger (reduction) and sympathy (for the company); and two behavioral, negative word-of-mouth (reduction) and purchase intention.

#### Procedures

The control and experimental groups read the mock news article, “Two Plus Has Repeatedly Polluted Yuseong Stream.” The control group then responded to the survey questions online. The other six groups (230 subjects) were next given their respective public apologies. The experimental groups then responded to the survey questions.

#### Experiment 2

##### Participants

A total of 2,800 people who live in Seoul, Korea between the ages of 20 to 59 (average 39.2) participated in the online survey. Among them, 600 participants were the control group and the other 2200 were allocated into 22 experimental groups. The survey was conducted between December 6^th^, 2012 and July 3^rd^, 2013, and implemented by a major professional research company in Korea. Participants were randomly selected from the company’s research pool (total population of 1.61 million), and they were rewarded with bonus mileage points for participating.

##### Materials

We developed a mock news story about a chemical company that was accused of discharging untreated wastewater. We left the specific names of the company and CEO in the mock news and public apologies blank (a) to minimize potential participant bias, and (b) to seem more realistic rather than using fake names.

Twenty-two types of public apology messages were developed as follows. First, we developed the baseline or basic public apology, which was then included in all apologies tested. Second, we added five different public apology messages — defense, acceptance (of responsibility), recovery, shame, and contact — to the basic apology. Third, using different combinations of two out of the five additional messages, we created ten composite public apologies (composite type I). Fourth, we combined three active messages — acceptance of responsibility (which includes willingness to pay liabilities), recovery, contact — and all five single messages (composite type II). Lastly, with the *recovery* statement, which does not include any money compensation message, we developed three different public apologies using different amounts of money compensation, denoted M1-M3. To measure participant responses to the news report of the crisis and subsequent apologies, we again used the effectiveness measures employed in Experiment 1 (Coombs & Holladay, 2008).

##### Procedures

The control and experimental groups read the same mock news report of the crisis event, and the experimental groups then read an apology prior to filling out the questionnaire. Each experimental group (22 total, 100 people each) was exposed to one of the 22 public apologies.

#### Experiment 3

The experiment was conducted via an online survey with 219 participants (111 males, 108 females). We selected five crisis types from those identified from previous studies where organizations have strong responsibility (see Experiment 1 Materials) (Coombs, 2014; Coombs & Holladay, 2002), and developed five short crisis event articles for each crisis type.

For each short article, we developed two types of public apology using the causal attributions: IC vs. EU. Each apology had four components: (1) the same regret message for both types, (2) one of the external or internal attributions; (3) one of the uncontrollable or controllable attributions — maintaining the same IC and EU pairings; and (4) the same *forbearance* from reoccurrence closing statement (see Table 5 for example). We presented the crisis situations to the participants in random order and then asked them seven questions (in random order) about the crisis situations, using a Likert-type seven-point scale. The seven effectiveness categories and the actual survey questions used for each are listed in Table 6 (Coombs & Holladay, 2008).

